# Ipsilateral stimulation shows somatotopy of thumb and shoulder auricular points on the left primary somatosensory cortex using high-density fNIRS

**DOI:** 10.1101/2024.09.16.612477

**Authors:** Ernest C. Okorie, Hendrik Santosa, Benedict J. Alter, Jacques E. Chelly, Keith M. Vogt, Theodore J. Huppert

## Abstract

**Significance:** Auriculotherapy is a technique based on stimulation applied to specific ear points. Its mechanism of active and clinical efficacy remain to be established. This study aims to assess the role that primary somatosensory cortex may play to validate auriculotherapy mechanisms.

**Aim:** This study examined whether tactile stimulation at specific auricular points is correlated with distinct cortical activation in the primary somatosensory cortex.

**Approach:** Seventeen healthy adults participated in the study. Tactile stimuli were delivered to the thumb, shoulder, and skin master points on the ear using von Frey filaments. Functional near-infrared spectroscopy was used to measure and spatially map cortical responses.

**Results:** This study revealed distinct hemodynamic activity patterns in response to ear point stimulation, consistent with the classic homunculus model of somatotopic organization. Ipsilateral stimulation showed specific cortical activations for the thumb and shoulder points, while contralateral stimulation showed less significant activity. Functional near-infrared spectroscopy effectively captured localized cortical responses to ear tactile stimuli, supporting the somatotopic mapping hypothesis.

**Conclusion:** These findings enhance the understanding of sensory processing with auricular stimulation and supports the concepts of auricular cartography that underpins some schools of auriculotherapy practice. Future research should explore bilateral cortical mapping and the integration of other neuroimaging techniques.

## 1 Introduction

Auriculotherapy is a therapeutic approach based on stimuli applied to the ear. Though non-specific electrical stimulation can be used, the most common method of auriculotherapy involves stimulation of specific points within an established cartography ^1^, which are believed to specific anatomic part or functions ^2^. Clinically, it is used to treat a wide variety of symptoms or conditions including postoperative pain, acute pain, chronic pain ^3^, anxiety ^4^, depression ^5^ and dementia ^6^. However, the physiological basis and clinical efficacy of auriculotherapy is still controversial and debated ^7^. Early evidence supporting this somatotopy came from the use of auriculotherapy for the treatment of lower back pain ^8^. More recent evidence provided by two studies ^9, 10^ confirmed the location of the point of thumb in the ear. Moura and Chaves demonstrated that auriculotherapy reduced chronic lower back pain ^11^. In contrast, a particular type of auriculotherapy, battlefield acupuncture, was not effective in reducing postoperative pain following tonsillectomy ^12^. In a meta-review of studies using auriculotherapy ^11^, 80% of studies (12 of 15) reported decreases in pain and the remaining 20% found no change relative to the control group. Of these 15 studies, only 7 studies ^13–19^ provided quantitative rating scores that could be used in the meta-analysis. Analysis of the 147 total patients and 159 controls from these studies revealed a reduction in subject reported pain rating (0-10 scale) of approximately −0.56 [95% CI = [−1.09 to −0.03]).

### 1.1 Primary somatosensory cortex somatotopy

The cortical homunculus, a common somatotopic map of the primary motor and primary somatosensory cortices, shows how the motor ^20^ or sensory stimulation ^21, 22^ of a body part activates a specific area of the cortex. A number of brain atlases have been developed that aim to break down the primary somatosensory cortex into even smaller regions, such as the Connectome Mapper ^23^, the study by Glasser and colleagues ^24^, and the Yale Brain Atlas ^25^. There have been past studies, mainly using fMRI, involving both motor tasks ^26, 27^ and tactile stimuli ^28^ showing the accuracy of the mapping, with specific activations shown in the lateral-medial and anterior-posterior directions of these regions. As many of these studies have looked at differentiating the digits of fingers and their somatotopic representation along the sensory ^29–32^ and motor cortices ^33, 34^,

### 1.2 Somatotopy of Auricular stimulation

We found two studies ^9, 10^ that looked at brain activity localization based on stimulation points on the ear, based on the Nogier system ^35^. While both previous studies focused on thumb point stimulation of the ear using fMRI, with the latter also stimulating the brain stem auricular acupoint, our study expands this scope by including shoulder auricular point stimulation and using fNIRS to overcome the inherent limitations of stimulation of the ear within an MRI head coil. A key limitation of the first study was its focus on a single stimulation point, while the second study, although it included an additional brain stem auricular acupoint, did not include points for anatomically distant body parts. In the present study, we examined the theory that there is a topographic map of acupuncture points on the ear to the same cortical region on the primary somatosensory cortex (S1) as the specific body parts ^36–38^, specifically the shoulder and thumb points ^35^. Two previous fMRI studies ^9, 10^ have suggested that this mapping exists, as evidenced by evoked brain changes observed in regions of S1 corresponding to the actual thumb, contralaterally in 6 subjects and bilaterally in 3 subjects ^9^, and in the secondary somatosensory cortex bilaterally when stimulating the thumb point and in the limbic and cortical areas of the pain matrix, with stronger activity ipsilaterally, when stimulating the brainstem point ^10^ in six subjects. Our study addresses these limitations by stimulating points for two body parts that are far apart anatomically and distant enough within the auricular cartography to enable distinct discrete stimulation of separate points.

### 1.3 Functional near-infrared spectroscopy

Functional near-infrared spectroscopy (fNIRS) is a noninvasive neuroimaging technique that uses low levels of light in the range of approximately 650-900 nm to measure changes in blood oxygenation in the brain. This technique was first described by Jobsis in 1977 ^39^ and has since been used in hundreds of brain imaging studies ^40^. Compared to other imaging methods, such as functional magnetic resonance imaging (fMRI), fNIRS is more portable and less expensive and does not require a complex or dedicated environment for imaging. FNIRS measurements are recorded through a customized head cap worn by the participant, where a set of discrete light source and detector components (optodes) are positioned spatially above the region of interest of the brain. Light from these source optodes is transmitted into the underlying tissue, where it diffuses and is optically scattered many times. Some of the light exiting the tissue is captured by fNIRS detector optodes placed a few centimeters away from the source on the surface of the scalp. These measurement pairs between discrete source and detector positions on the scalp record changes in the light transmitted through the diffuse volume of tissue between the source-detector pair, which can reach approximately 5-8 mm into the cortex of the brain at a typical source-detector separation of approximately 35 mm. By using a grid of these optodes, as shown in Figure 1, fNIRS can estimate the spatial changes in oxy- and deoxyhemoglobin associated with vascular and oxygen metabolic changes in the brain during activity.

**Figure 1.**
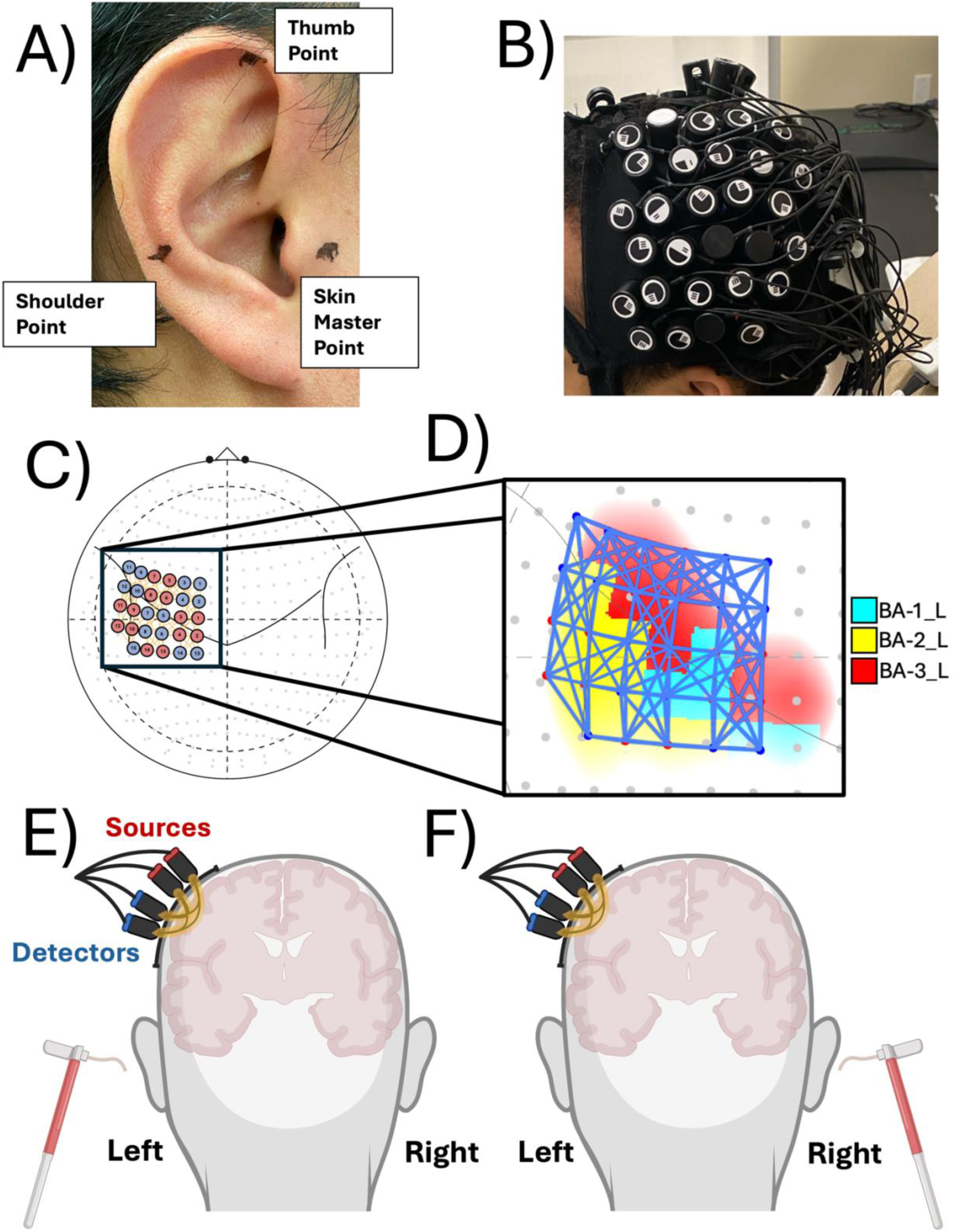
Functional near-infrared spectroscopy schematics and experimental setup A) Auricular points, based on the cartography of Nogier (Oleson & Kroening 1983, Nogier 1972), on the ear that were stimulated. B) Probe cap with sources and detectors on the subject’s head. C) High-density layout of the sources, in red, and the detectors, in blue. D) Probe layout over Brodmann areas 1, 2, and 3 over the left primary somatosensory cortex. The blue dots indicate the detectors, the red dots indicate the sources, and the blue lines indicate the channels. E) Schematic of the ipsilateral stimulation, where the left ear is stimulated and the probe is on the left hemisphere. F) Schematic of the contralateral stimulation, where the right ear is stimulated and the probe is on the left hemisphere. Bottom panels created using Biorender.com, photographs are of the authors.

The advantages of fNIRS over fMRI and EEG include its insensitivity to electrical artifacts ^41^, compatibility with free movement ^42^, and ability to be used in various environments ^43^. Moreover, fNIRS equipment is silent ^44^ and nonmagnetic. The portability and relative ease of setup of fNIRS systems facilitate a more realistic experimental environment, reducing participant discomfort and potential interference with the tactile stimulus response. The fNIRS cap also allows much greater access to the head, compared to having a subject within a head coil and in the center of an MRI scanner bore. These advantages make fNIRS a suitable candidate for this study which sought to investigate activated brain regions, in response to sensory stimulation, with greater precision and fewer methodological compromises.

## 2 Methods

### 2.1 Participants

The study involved seventeen healthy adults, comprising three females and fourteen males. Three participants were left-handed, and fourteen participants were right-handed, as determined through self-reports. Participants were scheduled for a single visit lasting approximately two hours. For their time and participation, they received financial compensation. Informed consent was obtained from each individual, in line with the ethical standards and guidelines approved by the University of Pittsburgh institutional review board (IRB).

### 2.2 Stimulation protocol

Tactile stimulation of the ear points was done using von Frey filaments ^22, 45, 46^. The stimulation sites (thumb, shoulder, and skin master points) were on the skin of the external ear, in accordance with the Nogier auriculotherapy point system ^35, 47^ as shown in Figure 1. Tactile stimulation was delivered using 26 g von Frey filaments. The von Frey filaments are nylon filaments that, when applied perpendicularly to the skin, exert a calibrated pressure on the skin. Von Frey filaments were then repeatedly applied to the test site ^22, 45, 46^. Stimulation for touch sensation was not overly painful and did not damage the skin. Safeguards to ensure this were established as part of the standardized operating procedures for these tests. First, the experimenters monitored verbal and nonverbal feedback from the participants. Participants were instructed multiple times to notify the experimentalist if the stimulation was intolerable. Finally, the participants could withdraw and stop the stimulation at any time. All of these safeguards minimized the risk to participants from touch stimulation.

### 2.3 Experimental design

To investigate the differential cortical responses within the left S1 to stimuli on both the ipsilateral ear and the contralateral ear, we developed the following ear stimulation paradigm. Participants were seated in a chair with a 45-degree semi-reclined position. Their legs were placed horizontally on a leg rest, ensuring a comfortable and stable position throughout the experiment. All sessions were conducted in a quiet room with participants having their eyes closed to minimize external visual distractions.

Tactile stimulation was presented with a blocked design timing. For each scan, 26 g von Frey filaments were used to stimulate the thumb point for 10 seconds, followed by a 15 second rest period, the shoulder joint point for 10 seconds, another 15 second rest period, and finally, the skin master point ^35^ for 10 seconds. This sequence was repeated five times throughout the scan, and three scans were performed. The first two scans targeted the ear contralateral to the fNIRS probe (right ear), and the final scan targeted the ipsilateral ear (left ear), as shown in Figure 1.

### 2.4 Self-ratings of pain and pressure levels

As a part of the prompt before the experiment, the sensation of pain was described as “any uncomfortable, distressing, or sharp feeling” induced by the filament. In contrast, pressure sensation was characterized as the “amount of force or weight perceived during the application of the filament, devoid of any painful quality”. After the conclusion of the experiment, participants were asked to rate the pain and pressure sensation from the filament on a scale of 0-10, where 0 represented “no sensation”, and 10 represented “the worst you’ve ever felt”.

### 2.5 Data collection and analysis

The fNIRS data were acquired using a NIRScout-2 (NIRx, GmbH, Berlin, Germany) continuous wave fNIRS system. We employed a high-density probe configuration to optimize left somatosensory cortex coverage and signal quality. As shown in Figure 1, the fNIRS probe array comprised 107 measurement channels, with 105 channels designated for long-distance measurements and two channels designated for short-distance measurements. The high-density probe was positioned on the left side of each subject’s head. Probe was aligned with the sagittal center line of the head and positioned approximately 3 cm off midline with source-1 aligned with Cz. Consequently, stimulation of the left ear was termed “ipsilateral stimulation,” aligning with the probe’s location, whereas stimulation of the right ear was referred to as “contralateral stimulation”. The spatial arrangement of the optodes was designed to facilitate a comprehensive examination of the cortical areas of interest. Specifically, the sources and detectors were placed such that the inter-optode distances were maintained at 15 to 44 mm for long-distance channels ^48, 49^. This configuration allowed long separation channels to probe deeper layers of cortical activity, thereby capturing neuronal hemodynamic responses and not scalp hemodynamics. Channel distances less than 15 mm were not included in the analysis.

Analysis was conducted to identify significant changes in cortical activation, as measured by changes in the concentrations of oxygenated hemoglobin (HbO) and deoxygenated hemoglobin (HbR) in response to tactile stimuli. Preprocessing and analysis were performed using MATLAB (Version 2020a) and the AnalyzIR Toolbox ^50^. The grid layout of the channels, boxplots and histogram were generated using Python (version 3.12.00). We first converted the raw fNIRS data to optical density ^51^ and converted that to relative ΔHbO and ΔHbR concentrations using the modified Beer–Lambert law ^52, 53^. A baseline principal component analysis filter was applied, as recommended by Franceschini et al. ^54^, where eigenvectors of the baseline were removed until 80% of the spatial covariance was explained. Finally, the data were resampled to 4 Hz. The group data were analyzed using a general linear model (GLM) ^55, 56^ approach tailored for block design studies.

Statistical analysis of the fNIRS data was done using the AnalyzIR Toolbox ^50^, which was developed by our group. A robust, pre-whitening, general linear model ^57^ was used for the first-level analysis to provide estimates of the evoked hemodynamic responses for the three task conditions (thumb, back, and skin master). In the second-level analyses, we used a weighted, robust, linear mixed-effects model. The estimate of the parameter covariance from the first-level linear model was used to pre-whiten and weight the data in this group analysis. The linear mixed effects model was then solved iteratively using a bisquare robust estimator to decrease the influence of outlier data points. The participant-reported pain ratings were used as an additional covariate of interest in the model. Following the mixed effects analysis, regions of interest (ROI) averages were computed. Regions were identified based on probe sensitivity, specifically within the left hemisphere Brodmann areas BA1, BA2, and BA3, which correspond to the primary somatosensory cortex, utilizing the Colin27 atlas included ^58^ in the analysis toolbox ^59^.

Statistical significance for the estimated evoked brain activity was identified based on a Benjamini Hochburg ^60^ false discovery corrected threshold of q<0.05. The resultant channel maps are illustrated as heatmaps, plotting the source number on the horizontal-axis and the detector number on the vertical-axis, highlighting only cells meeting these statistical criteria. The arrangement of source numbers on the heatmap is based on the MNI x-coordinates taken from the toolbox, ordered from the furthest to closest relative to the origin. Similarly, the detector numbers follow the same ordering principle. Consequently, the heatmap’s bottom left corner displays more lateral channels, while the top right corner indicates more medial channels according to this spatial organization.

## 3 Results

### 3.1 Individual auricular point hemodynamic responses versus baseline

The respective channel mapping and statistical analysis for each stimulation condition—thumb, shoulder, and skin master—are depicted in Figure 2 for the ipsilateral and contralateral stimulations.

**Figure 2.**
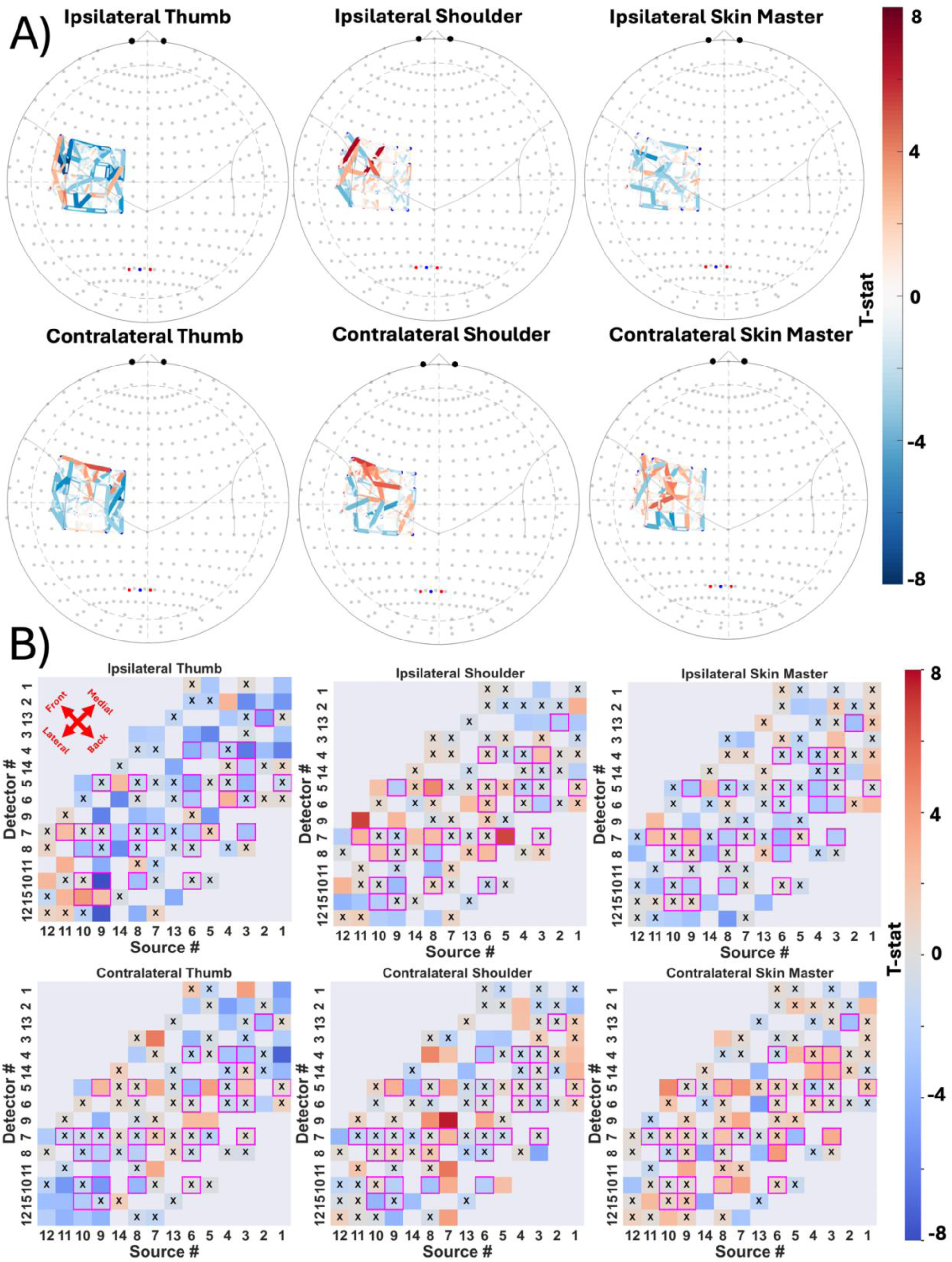
Hemodynamic response to ipsilateral and contralateral ear stimulation of the three auricular points shown on a topographic map of the head and a grid layout of the channels showing a color-coordinated t-statistic map. Gray cells indicate that the source-detector pair does not have a channel between them, a cell with an “X” represents that the absolute value of the t-statistic of the channel is less than 2, and cells with a pink border show that the channel is within the region of interest. A) Topographic map. B) Grid layout.

A total of 32 distinct channels within the specified ROIs met the inclusion criteria. The channels that surpassed the statistical threshold of q<0.05 are shown in Figure 2.

For ipsilateral stimulation, the results varied by stimulation point. Specifically, for the thumb point, four channels exhibited a significant positive t-statistic, while twelve channels had a significant negative t-statistic. Shoulder point stimulation resulted in seven significant positive channels and six significant negative channels. Finally, the skin master point showed similar results, with two significant positive channels and eleven significant negative channels.

In contralateral stimulation, the patterns were slightly different. The thumb point had five significant positive channels and six significant negative channels. The shoulder point revealed two significant positive channels and five significant negative channels. The skin master point had seven significant positive channels and two negative channels.

### 3.2 Contrast between the thumb and shoulder auricular points and spatial differences

Figure 3 displays the channel map and the heatmap for all channels, with blue highlights showing the channels within the regions of interest (ROIs) for the contrasts between thumb and shoulder stimulations under both the ipsilateral and contralateral conditions. Specifically, the map shows where the hemodynamic response to thumb stimulation was greater than that to shoulder stimulation and vice versa. Finally, the heatmap in Figure 4 encompasses only those channels within the ROIs for both ipsilateral and contralateral conditions as well as their t-statistic values, clearly depicting where the thumb versus shoulder and shoulder versus thumb hemodynamic responses differ.

**Figure 3.**
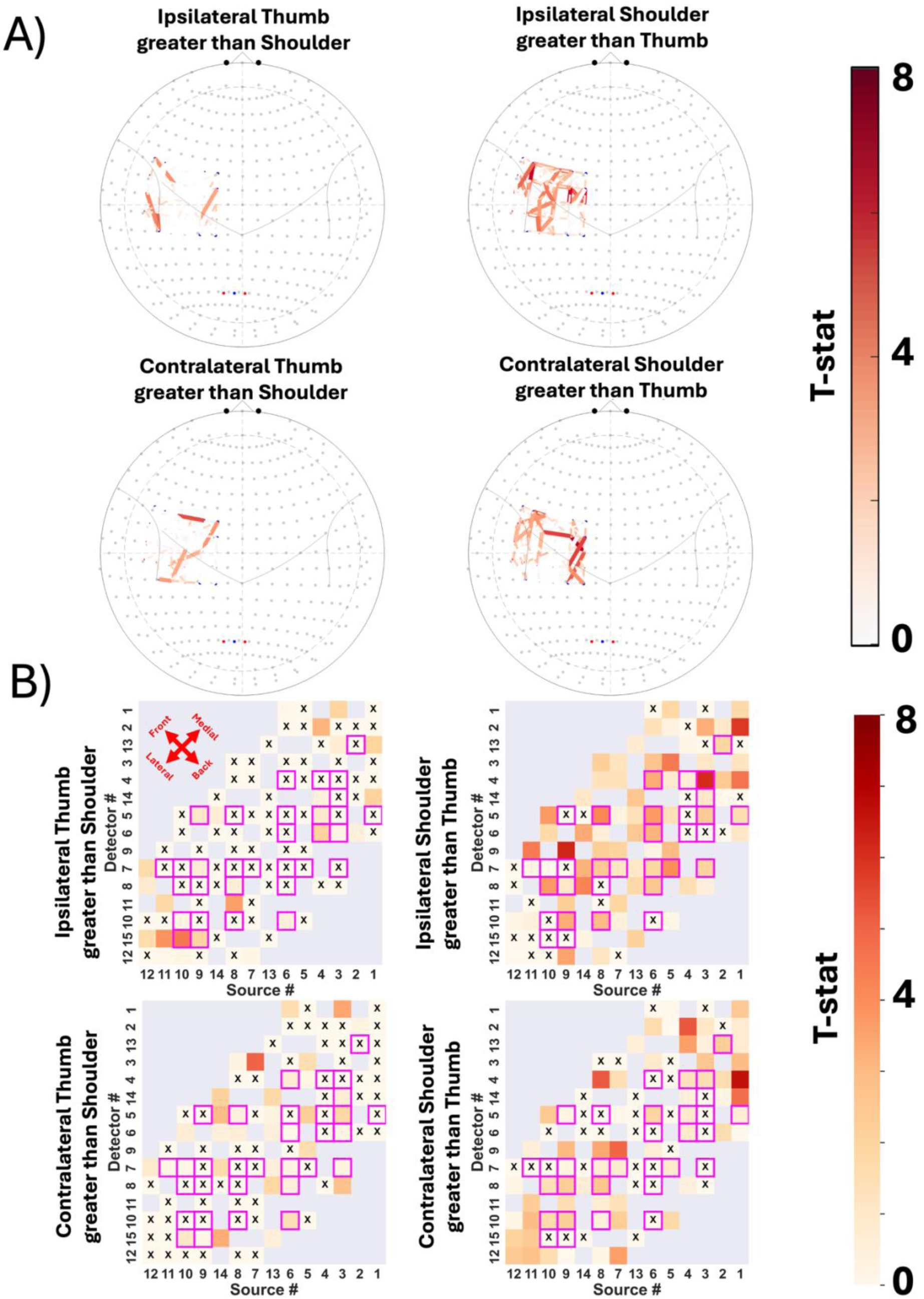
Ipsilateral and contralateral thumb greater than shoulder or shoulder greater than thumb contrasts shown on a topographic map of the head and a grid layout of the channels showing a color-coordinated t-statistic map. Gray cells indicate that the source-detector pair does not have a channel between them, a cell with an “X” represents that the value of the t-statistic of the channel is less than 0, and cells with a pink border show that the channel is within the region of interest. A) Topographic map B) Grid layout.

**Figure 4.**
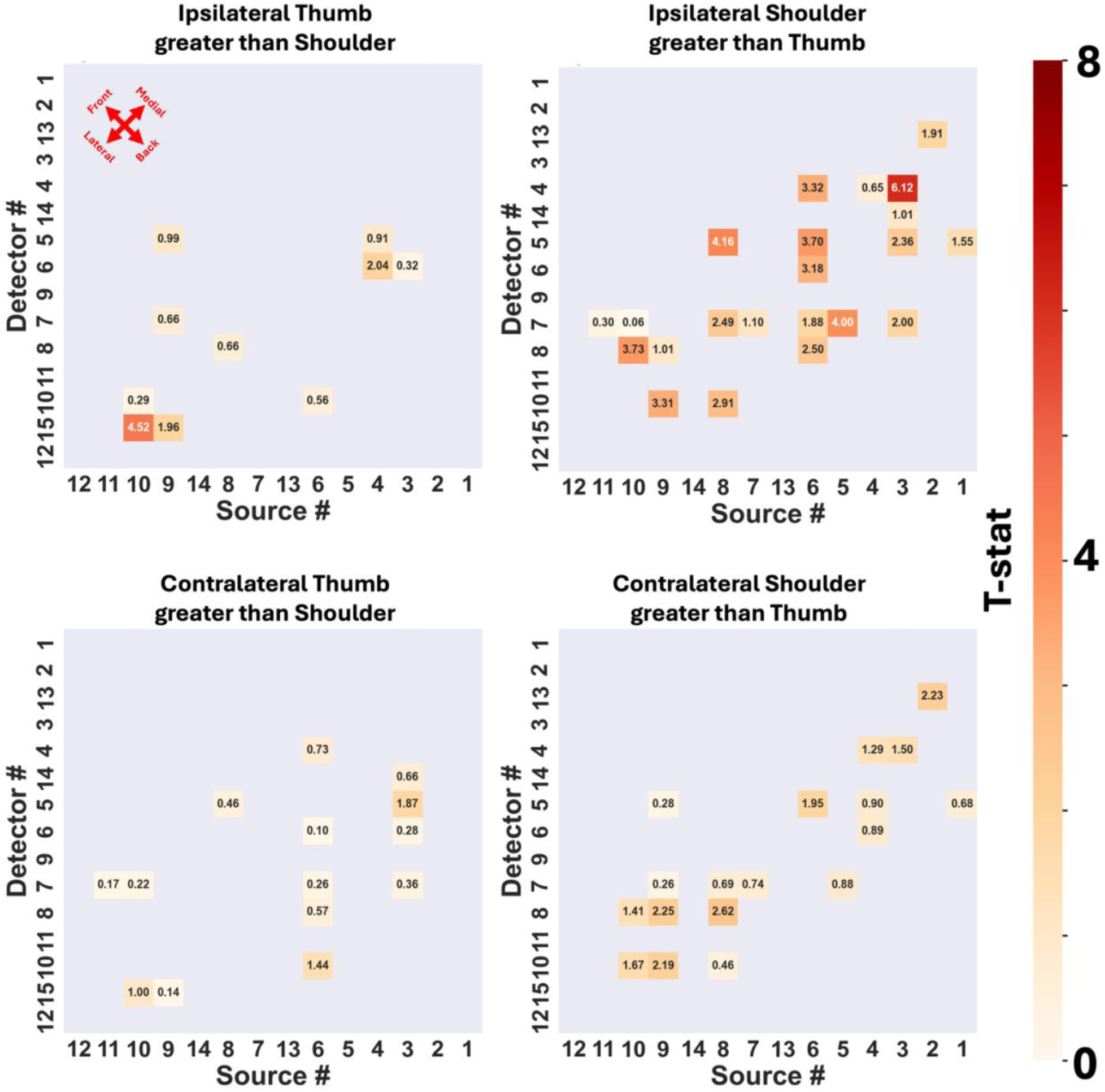
Ipsilateral and contralateral either thumb greater than shoulder or shoulder greater than thumb contrasts with only the channels that are within the regions of interests also showing the t-statistic value.

Stimulation location contrasts showed varying levels of significance across different channels. For the ipsilateral thumb greater than shoulder contrast, the highest T-statistic was observed for the channel between the source-detector pair S10-D15 (T = 4.5, Q-value = 3.6E-4). Conversely, the ipsilateral shoulder greater than thumb contrast revealed its peak t-statistic for channel S3-D4 (T = 6.1, Q-value = 5.6E-7). In the case of contralateral contrasts, the thumb greater than shoulder comparison yielded the highest T-statistic for S3-D5 (T= 1.9, Q-value = 1.7E-1), while the shoulder greater than thumb contrast showed a maximum T-statistic for channel S8-D8 (T = 2.6, Q-value = 4.7E-2). Notably, the spatial distance between channels S10-D15 and S3-D4 is 63.6 mm, and that between channels S3-D5 and S8-D8 is 28.8 mm. Table 1 shows the statistical values of each of these 4 selected channels.

**Table 1.** The channels selected based on having the highest t-statistic values from each contrast condition and these channels’ statistical values within the ROI.

| Side | Condition | Source # | Detector # | Beta | Standard Error | T-statistic | Q-value |
| --- | --- | --- | --- | --- | --- | --- | --- |
| Ipsilateral | Thumb-Shoulder contrast | 10 | 15 | 3.96 | 0.88 | 4.52 | 0.00036 |
| Ipsilateral | Shoulder-Thumb contrast | 3 | 4 | 3.46 | 0.56 | 6.12 | 0.00000056 |
| Contralateral | Thumb-Shoulder contrast | 3 | 5 | 0.97 | 0.52 | 1.87 | 0.17 |
| Contralateral | Shoulder-Thumb contrast | 8 | 8 | 1.53 | 0.58 | 2.62 | 0.047 |

### 3.3 Self-reported pain and pressure levels

Figure 5 shows a boxplot comparing pain and pressure levels between ipsilateral and contralateral stimulations for each specific point. A histogram displaying the distribution and frequency of pain and pressure level ratings is included in Figure 5. Both ratings centered around 2-3 on the subjective numerical rating scales used.

**Figure 5.**
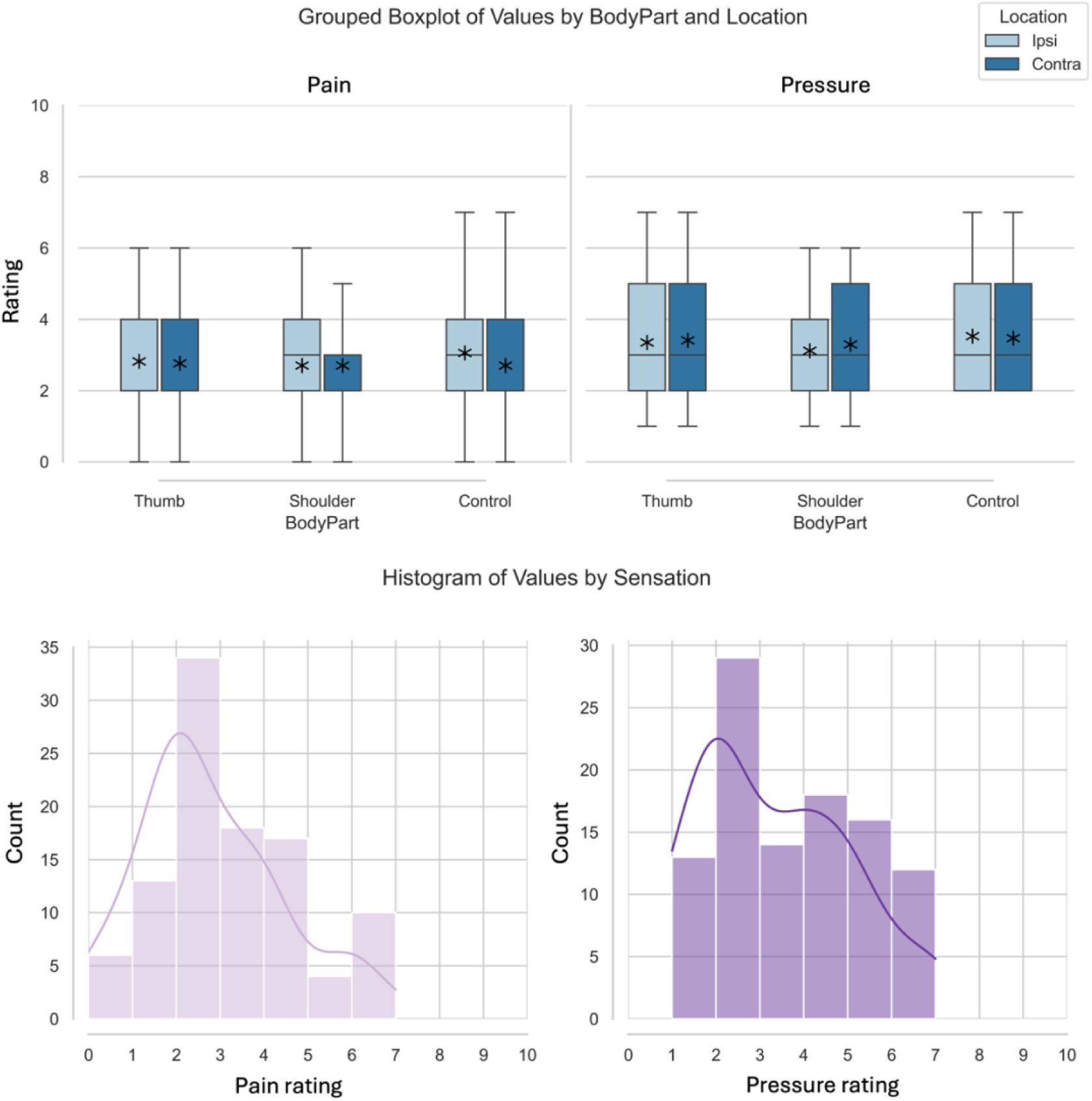
Boxplot and histogram of the self-reported pain and pressure levels from 0 to 10. The line inside the box represents the median of the data, the box itself shows the interquartile range, this range contains the middle 50% of the data. The whiskers extend to the furthest data point within 1.5 times the IQR from the quartiles. The stars represent the average. Legends: Ipsi = Ipsilateral, Contra = Contralateral

## 4 Discussion

### 4.1 Findings

This study demonstrated that von Frey filament stimulation of the left ear at the thumb vs, shoulder auricular points resulted in distinct localized hemodynamic activity reflecting somatotopic differences in left S1. Despite only stimulating subject’s ears, our group-level results demonstrate significant differences in brain responses within the primary somatosensory cortex. Specifically, ipsilateral thumb point stimulation resulted in responses primarily in lateral (S1). In contrast, shoulder auricular point stimulation was associated with more medial S1 hemodynamic activity. These patterns of responses to auricular stimulation support the concept of body parts represented with an auricular homunculus that maps somatotopically onto S1 with experimental stimulation of the ear. These findings provide additional support for the theoretical framework that underpins point-stimulation based auriculotherapy (i.e. with needles, pressure, or cryotherapy), distinct from less localized electrical stimulation of the ear (i.e. electro-auriculotherapy or trans-auricular vagal nerve stimulation).

### 4.2 Strengths and Limitations

A limitation when using fNIRS is the variability in cranial anatomy among the subjects, which can influence the positioning of probes and, as a result, the detection of signals ^61, 62^. Although the standard probe layout provides a general framework for optode placement, individual differences in cortical folding and size, and positioning of the array may lead to variations in data acquisition that are not easily controlled without personalized anatomical mapping. The depth of penetration of fNIRS, which is limited to approximately 2-3 cm on the adult head ^63, 64^, poses a challenge for detecting deeper hemodynamic activity ^65, 66^. This limitation may result in an underestimation of deeper hemodynamic changes and an overrepresentation of superficial activity. We also only had the probe located on the left hemisphere, which means we were not able to compare bilaterally whether these ipsilateral and contralateral stimulation sites held true on the right hemisphere. Additionally, it has lower temporal resolution than EEG ^67^ and lower spatial resolution than fMRI ^42, 68^.

### 4.3 Implications

By better understanding the localized brain responses to auricular stimulation, auriculotherapy could be more precisely studied and potentially improved as an effective treatment for chronic pain. This study contributes to a better understanding of how tactile stimulation of distinct auricular points specifically maps to representation within the ipsilateral primary somatosensory cortex. This finding supports the notion that the sensory system is highly organized and subtle variations in location of auricular stimuli can produce different brain responses.

## 6. Author contributions

EO contributed to the development of the manuscript, conducted data collection, and performed data analysis. HS contributed to data analysis and development of the manuscript. BJA contributed to the design of the study, interpretation of results, and the development of the manuscript. JEC contributed the design of the study, interpretation of results, and the development of the manuscript. KMV contributed to the design of the study, interpretation of results, and the development of the manuscript. TH contributed to the design of the study, oversaw data collection and analysis, contributed to interpretation of results and development of the manuscript.

## 7. Code and Data Availability

MATLAB and Python code files are available upon request.

## 8. Acknowledgments

The research reported in this publication was supported by K23NS123429.

## Author Biographies

**Ernest Okorie** is an undergraduate Natural Sciences major at the University of Pittsburgh, set to graduate in December 2024. He serves as a research assistant in the Huppert Lab and has also been a teaching assistant for General Chemistry, Organic Chemistry, and General Biology courses.

**Theodore J. Huppert**’s lab develops multimodal neuroimaging methods including MRI, MEG, EEG, diffuse optical imaging (NIRS), and PET imaging to provide a more complete modality independent picture of the brain through technology cross-validation, multimodal (statistical) data fusion, and underlying state-space modeling of the cerebral physiology. The specialty of his lab is near-infrared spectroscopy methods applied to unique neuroimaging scenarios (e.g., ambulatory movement, field-deployable, and pediatric brain imaging) and in concurrent multimodal experiments (e.g., NIRS/MEG and NIRS/EEG/fMRI).

